# CRISPR/Cas9-mediated mutagenesis of the *white-eye* gene in the tephritid pest *Bactrocera zonata*

**DOI:** 10.1101/2025.04.22.650062

**Authors:** Albert Nazarov, Tamir Partosh, Flavia Krsticevic, Dimitris Rallis, Yael Arien, Guy Ostrovsky, Reut Madar Kramer, Eyal Halon, Alfred M. Handler, Simon W. Baxter, Yoav Gazit, Kostas D. Mathiopoulos, Gur Pines, Philippos A. Papathanos

## Abstract

*Bactrocera zonata* is a highly invasive agricultural pest that causes extensive damage to fruit crops. The Sterile Insect Technique (SIT), a species-specific and environmentally friendly pest control method, depends on the availability of Genetic Sexing Strains (GSSs) to enable efficient mass production of males for sterile release. However, no GSS currently exists for *B. zonata* limiting SIT applications targeting this important invasive pest. Here, we report two key advancements toward GSS development in this species. First, we present a high-quality, chromosome-level genome assembly from male *B. zonata*, identifying two scaffolds derived from the Y chromosome, which represent potential targets for future male-specific genetic engineering. Second, we demonstrate the feasibility of CRISPR/Cas9 genome editing in *B. zonata* by generating stable, homozygous white-eye mutants through targeted disruption of the conserved *white-eye* gene. This visible, recessive phenotype serves as a proof-of-concept for developing selectable markers in this species. Together, these results provide foundational genomic and genetic tools to support the development of GSSs in *B. zonata*, advancing the potential for sustainable, genetics-based pest control strategies.

## Introduction

*Bactrocera zonata*, commonly known as the peach fruit fly, is a highly invasive and economically damaging agricultural pest. *B. zonata* is highly polyphagous, infesting almost 50 commercial and wild host plants, including peach, mango, guava, apricot, citrus, eggplant, tomato, apple and loquat (White and Elson-Harris 1992; Zingore et al. 2020). Native to South and Southeast Asia, it has now spread to more than 20 countries, including wide regions of Africa and the Middle East, since its initial establishment in Egypt in 1997 (Ni et al. 2012; De Meyer et al. 2007). *B. zonata* is a strong flier capable of dispersing 25 miles or greater in search of hosts, is highly adaptable to different habitats and has a relatively short generation time enabling multiple generations per year (Zingore et al. 2020). Isolated detections in European countries like France and Austria have raised concerns about its potential establishment in Southern Europe, where favorable climatic conditions and abundant host plants could facilitate its spread, resulting in its listing as an A1 quarantine pest in the European and Mediterranean Plant Protection Organization (EPPO) countries (EPPO Global Database 2023). The annual costs of damage in the Near East are estimated at 320 million EUR and in Egypt at 190 million EUR per year (El-Gendy 2022). Current pest management strategies predominantly rely on insecticides, yet these raise concerns about environmental impact and the evolution of insecticide resistance (Bourtzis & Vreysen, 2021). Therefore, there is a pressing need for alternative, environmentally friendly, and species-specific pest control strategies.

The Sterile Insect Technique (SIT) is a species-specific, environmentally-friendly approach for the area-wide integrated pest management (AW-IPM) of tephritid fruit flies. SIT involves mass-rearing, sex separation, sterilization, and subsequent release of sterile males into pest-infested areas. When wild females mate with these sterile males, they lay non-viable eggs, leading to suppression and potential eradication of the local pest population if releases are sustained. The success and cost-effectiveness of SIT programs significantly depend on the availability of genetic sexing strains (GSSs) that enable the mass-production and release of only sterile males (Augustinos et al. 2017; Dyck et al. 2021). Male-only releases are essential because sterile females can cause fruit damage through oviposition, reducing crop quality and consumer acceptance, and may also divert sterile males from mating with wild females, diminishing the effectiveness of the SIT. GSSs typically incorporate sex-specific selectable markers, such as mutations affecting pupal color (*white-pupae, black-pupae*), eye color (*white-eye, red-eye*), or temperature-sensitive lethality (*tsl*), linked to sex-determining loci on sex chromosomes (Franz, Bourtzis, and Cáceres 2021; Augustinos et al. 2017). In *Ceratitis capitata* (medfly), successful GSSs were generated through the translocation of selectable markers on the Y chromosome, a process that required complex genetic approaches, feasible only in species with well-developed genetic tools. In species with limited genetic resources, such as *B. zonata*, these classical genetic methods remain challenging due to the labor-intensive and time-consuming isolation of suitable mutants.

Two recent advances in CRISPR/Cas9 technology offer great promise in overcoming such hurdles in non-model species and accelerate the development of GSSs. Firstly, CRISPR/Cas9 genome-editing technologies offer a powerful alternative to classical mutagenesis, enabling precise targeting and modification of candidate genes associated with selectable phenotypes. Secondly, CRISPR/Cas9 homology-directed repair can mediate precise translocation (knock-in) of a selectable marker on the Y chromosome. However, the lack of high-quality genomic resources, particularly detailed genome assemblies and Y chromosome data, remains a critical bottleneck for developing such strains in non-model pests. To address these gaps, we present here two major advances toward the development of GSSs in *B. zonata*. First, we generated a highly contiguous, chromosome-level genome assembly of *B. zonata*, derived from male flies of a laboratory colony established in 2022 from field-caught flies in Israel. Through comparative genomics, we identified both autosomal and sex-linked scaffolds, including two scaffolds derived from the Y chromosome. These Y-linked scaffolds represent potential targets for future knock-in of selectable markers, a crucial step for enabling male-only production and advancing the development of effective GSSs.

Second, as a proof-of-concept for genetic manipulation in *B. zonata*, we successfully applied CRISPR/Cas9-mediated mutagenesis to disrupt the *white*-*eye* gene, which is essential for eye pigmentation (Ewart and Howells 1998). Mutations in the *white-eye* gene have been successfully established in other Tephritidae, including *C. capitata* (Meccariello et al. 2017), *Zeugodacus cucurbitae (Paulo et al. 2022), Bactrocera oleae (Meccariello et al. 2020), Bactrocera dorsalis (Zhao et al. 2019)* and *Bactrocera tryoni* (Choo et al. 2018). Consistent with these recent works, we continue the use of calling this gene *white-eye* instead of *white*, as it was originally named in *Drosophila*, to easily distinguish it from the common *white-pupae* mutation. Our successful generation of stable *white-eye* mutant lines demonstrates the feasibility of CRISPR-based genome editing in *B. zonata*, providing a robust foundation for future development of genetic sexing strains specifically tailored for SIT programs targeting this important pest.

## Materials and Methods

### Fly rearing

*Bactrocera zonata* flies were obtained from the Israel Cohen Institute for Biological Control. Fly colonies were maintained at 26 ± 1 °C, 65% ± 5 RH, under a 14/10 h light/dark cycle. Adults were fed a mixture of sucrose and hydrolyzed yeast (3:1) in water provided at lib. Larvae were fed on a diet containing bran (268 g/L), brewers yeast powder (81 g/L), sucrose (121 g/L), nipagin (2 g/L), HCl (25%, 16 mL/L), and water (510 ml/L). Larvae emerging from injected eggs (G0s) were reared on a modified diet of 50% larval diet mixed with 50% agarose solution. Vermiculite was added to the bottom of the rearing cage to allow pupation.

### *Bactrocera zonata* genome sequencing and assembly

*Bactrocera zonata* long-read genome assembly was performed using SRA-deposited genomic resources -Illumina: SRR29319676, SRR29319677; HiC: SRR29319678; Nanopore: SRR29299502, SRR29299503). Noisy Nanopore reads were error-corrected with FMLRC2 (Mak et al. 2023) using male Illumina reads and assembled using Flye (Kolmogorov et al. 2019). The resulting draft assembly was purged twice using purge-dups (Guan et al. 2020), followed by two rounds of polishing with male Illumina reads using Pilon (Walker et al. 2014). In the scaffolding step, reads from the Hi-C library were mapped to the polished contigs using the Arima-HiC mapping pipeline (ArimaGenomics 2023) and contigs were scaffolded using YAHS (Zhou, McCarthy, and Durbin 2023). The completeness of the assembly was assessed with BUSCO (Simão et al. 2015) using the diptera_odb10 database. Repetitive regions were identified and annotated using RepeatModeler2 (Flynn et al. 2020) and RepeatMasker (Tarailo-Graovac and Chen 2009). The final assembly and downstream scaffolding with HiC data produced a total of 1021 scaffolds with an N50 of 81 Mb, resulting in an assembled genome size of 672 Mb. The assessment of genome completeness using Diptera BUSCOs suggested a 99% completeness (Single copy; 97.6 %, Duplicated; 1.4 %, Fragmented; 0.3 %).

### Annotation and synteny analysis

The assembled *B. zonata* genome was annotated using Funannotate V1.8.15 (Palmer and Stajich 2020) with the GeneMark module and the following RNA-seq datasets (SRR4024785, SRR4024786, SRR4024787, SRR4024788). For the synteny analysis with *C. capitata*, the manually curated annotation of the Ccap_2.1 assembly (GCF_000347755.3) were mapped on the EGII_3.2.1 assembly (GCA_905071925.1) using minimap2 (Li 2018) and converted to annotations using bedtools (Quinlan and Hall 2010) and UCSC tools (Perez et al. 2025). Homologous transcript sequences between *B. zonata* and *C. capitata* were found with BLASTn (Perez et al. 2025) filtering for evalue < 1e-4 and percent identity > 80. The detection of collinear blocks of orthology between *C. capitata* and *B. zonata* was performed using MCScanX (Perez et al. 2025) and visualized with SynVisio (Venkat and Carl 2020) (Venkat and Carl 2020).

### Identifying sex chromosome derived scaffolds

R-CQ and KAMY (Rallis et al. 2023) were used to identify assembly regions derived from sex chromosomes, using male and female Illumina reads (SRR29319676, &female). For the R-CQ algorithm, reads were mapped to the *B. zonata* assembly using bwa-mem (Li and Durbin 2009). Single-mappers from male- and female-specific alignments were selected by removing reads with XA:Z and SA:Z tags from the alignment file, which was then converted to bedGraph using bedtools (Quinlan and Hall 2010) and then to BigWig with bedGraphToBigWig (Perez et al. 2025). The BigWig files from the mapping of Male and Female reads were used as input to R-CQ. Scaffold 1 was used to normalize sequencing depth between male and female libraries and the male/female ratio of depth was calculated from the mean of per-base coverage in sequence windows of 50 bp. The calculated male/female depth-ratio values were plotted for every respective sequence window with karyoplotR.

### Coverage-based mapping and PCR verification of Y chromosome scaffolds

To evaluate Y-linkage of two putative Y-derived scaffolds (scaffolds 10 and 16), we mapped male and female WGS reads to both scaffolds. Reads from each sex were aligned to the genome using bowtie (Langmead et al. 2009) with parameters set to report all alignments (-a) and allow no mismatches (-v). Per-base coverage for each sex was computed using the genomeCoverageBed sub-command from bedtools, option -d (Quinlan and Hall 2010). The male and female per-base coverage counts were normalized by total library size and joined. To visualize sex-specific differences in coverage, mean coverage was calculated over 5 kb sliding windows, log2-transformed, and plotted in ggplot2 with female values displayed on the negative axis for ease of comparison. For the *MoY* region specifically, mean coverage was computed over 50 bp windows. To validate the Y-specificity of these scaffolds, PCR primers were designed at regular intervals across their entire length, ensuring no predicted off-target amplification in the rest of the genome assembly. The 13 copies of *MoY* were identified via tBLASTn (Altschul et al. 1990) searches and manually annotated. Predicted *BzMoY* coding sequences, proteins, and flanking regions were aligned to generate a consensus sequence and identify copy-specific or shared single nucleotide polymorphisms (SNPs).

### *B. zonata white-eye* gene analyses

Reciprocal tBLASTn searches of the *white-eye* from *B. tryoni* identified a single, high-likelihood ortholog on scaffold 3 of the *B. zonata* assembly (coordinates 34,350,949 - 34,370,584), which was manually re-annotated to correctly describe the gene model. Sequences of Tephritidae White-eye proteins along with the 2 Mbp of surrounding sequence were downloaded from NCBI: *B. tryoni* (LOC120776067 in GCF_016617805.1), *B. dorsalis* (LOC105224216 in GCF_023373825.1), *B. latifrons* (LOC108974879 and GCF_001853355.1), *B. neohumeralis* (LOC126758950 in GCF_024586455.1), *Bactrocera oleae* (106620505 in GCF_042242935.1*) C. capitata* (101458180 in GCF_000347755.3), *A. ludens* (LOC128858475 in [GCF_028408465.1](https://www.ncbi.nlm.nih.gov/assembly/GCF_027943255.1/)), *A. obliqua* (LOC129239220 in GCF_027943255.1). Multiple fragments of the *white-eye* gene were amplified and Sanger sequenced from multiple individuals separately to confirm the assembly and identify any polymorphisms in our laboratory strain (Primers list in **Table 1**).

### CRISPR/Cas9 reagent

gRNAs targeting the *B. zonata white-eye* gene were selected using Geneious Prime (2023.1.2) and the *B. zonata* genome assembly as a reference for off-targets. We selected gRNA-1 (CAGGAGGTGCTAATAAGAGGCGG) matching no other site with off-targets for synthesis (IDT). The gRNA was reconstituted in an injection buffer (5 mM KCl and 0.1 mM phosphate buffer, pH 6.8) to a final concentration of 100 μM. 1 µl gRNA was mixed with 1 µl tracer RNA and incubated at 90°C for 5 minutes, then at 25°C for 5 minutes. The mixture was then complexed with 2 µl Cas9 protein (PNA Bio, 1000 ng/µl) by incubating at room temperature for 5 minutes, followed by adding 1 µl of 5x injection buffer to complete the injection mix to a total volume of 5 µl.

### Egg collection and microinjection

To induce vigorous egg laying, we used a fruit facsimile made from cleaned plastic 100 ml yogurt bottles (e.g. Actimel) pierced with a sterile needle to create small holes for females to lay eggs (Gazit and Akiva 2017). A similarly perforated 50 ml Falcon tube, containing freshly cut citrus fruits, such as kumquats or oranges, was placed inside the plastic bottle, to elicit olfactory-driven behaviors. Females were given a two-hour window to oviposit eggs. Following this period, eggs were transferred to collection baskets using distilled water. Eggs were treated with a 0.5% sodium hypochlorite for 30 seconds to soften their chorion, followed by rinsing under running water for one minute.

Eggs were aligned on a transparent 0.9% agarose step to secure them during microinjection. The agarose step was produced by pouring molten agarose into a 35mm Petri-dish containing a glass microscope slide in its center as a mold and allowing it to solidify for 20 minutes. Once solidified, the agarose puck was carefully removed from the petri-dish and inverted to release the slide, leaving its negative impression. Using a clean blade, the agarose puck was trimmed parallel to the slide and cut into small square pieces each containing two faces (planar surfaces) offset by 1mm. Eggs were directionally aligned in series against the 1mm vertical riser between the two agarose faces and left on the agarose until hatching. Microinjections were performed between 2–4 hours after egg laying (AEL) using quartz glass capillaries (1 mm outer diameter, 0.70 mm inner diameter) pulled with a Sutter

P-2000 Laser Puller (settings: Heat 750, Filament 4, Pull 165, Velocity 40, Delay 150). Injections were performed using a Eppendorf FemtoJet 4I injector and a Narishige MM-94 micromanipulator mounted on a Olympus IX53 inverted microscope. Following injection, eggs were incubated at 26°C for 48 hours. Hatched G0 larvae were transferred to 12-well plates containing G0 larval food, which differed from standard larval food by the addition of a 0.5% agarose solution in a 1:1 ratio to maintain moisture and solidify the diet, with each well accommodating up to 5 larvae. These plates were placed in a box with 0.5 cm of ground vermiculite to collect pupae at the end of larval development.

### Live genotyping of mutants

For live genotyping, one mesothoracic leg was removed from each CO2 anesthetized flies. The leg was carefully excised with sterile scissors and forceps under a dissecting microscope. DNA was extracted using the Chelex protocol for DNA extraction (Musapa et al. 2013). PCR amplification of the first exon of the *white-eye* containing the gRNA1 target site was performed using primers PrimerD_white_ex1 and PrimerF_white_ex1, and amplicons were Sanger sequenced.

### Microscopy

Adult flies were placed at -20°C for one hour, followed by a 10-minute thawing period to allow evaporation of excess moisture. Specimens were mounted vertically using clay putty to facilitate imaging of the facial features, or laterally for full body imaging. Images were captured using a Nikon SMZ25 stereomicroscope equipped with a DS-Ri2 camera (NIKON Corporation, Japan). NIS-Elements software (ver 5.41.01) (NIKON) was used to capture images. Acquisition and z-stacking were captured by the Z-Series Auto option to increase depth of field.

## Results

As part of an international collaboration coordinated by the International Atomic Energy Agency, we generated new genomic data for five members of the Tephritid family - *Anastrepha fraterculus, An. ludens, B. dorsalis, B. zonata* and *Zeugodacus cucurbitae* (BioProject: PRJNA1082643; H. Djambazian in preparation). For *B. zonata* we combined Oxford Nanopore long-read whole-genome sequencing (WGS) of male genomic DNA, with separate Illumina WGS of both sexes, and male Hi-C data. After error correction and *de novo* assembly, the final assembly comprised 1021 scaffolds with a total size of 672 Mb, an N50 of 81 Mb, and 99% BUSCO gene completeness. Notably, the ten largest scaffolds ranged from 7 to 102 Mb, underscoring the highly contiguous nature of the assembly.

Synteny was assessed by mapping collinear blocks of predicted genes between *B. zonata* and the chromosome-level Eg-II assembly of *C. capitata* (medfly). Our analysis revealed that the seven longest *B. zonata* scaffolds corresponded to the typical five autosomes found in Tephritids (**Fig. 1A**, (Yesmin et al. 2019). Specifically, medfly chromosome 2 aligned with *B. zonata* scaffolds 4 and 7; chromosome 3 to scaffolds 6 and 5; chromosome 4 to scaffold 1; chromosome 5 to scaffold 3; and chromosome 6 to scaffold 2, respectively. These results suggested that several scaffolds likely represent complete chromosomes or full chromosomal arms.

**Figure 1.**
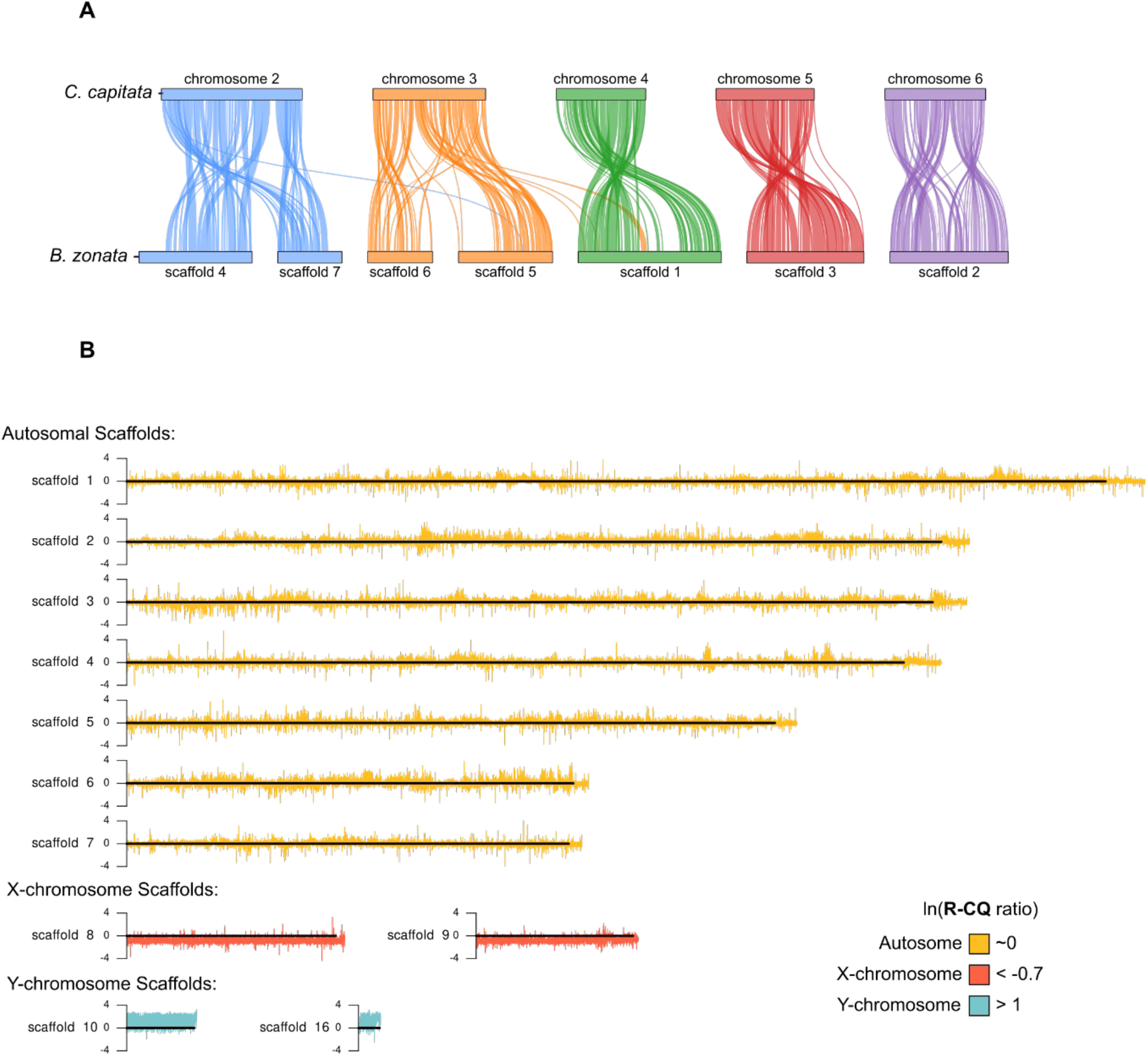
Chromosome-scale genome assembly and identification of sex chromosomes in *Bactrocera zonata*. **A)** Synteny analysis between *B. zonata* (bottom) and *Ceratitis capitata* (top) shows conserved collinear blocks across the seven largest scaffolds, corresponding to the five autosomal pairs in Tephritidae. Colored vertical ribbons connect blocks of syntenic orthologous genes across the two species. Each color represents a distinct *C. capitata* chromosome. **B)** Identification of sex-linked scaffolds using male/female coverage ratios. Karyoplot of R-CQ ratio of the 7 autosomal and 4 sex chromosome scaffolds in the *B. zonata* assembly. lnMedian(R-CQ) ratio of ∼0 is expected for autosomal sequences, >1 indicates Y-derived sequences and < - 4 indicates X-derived sequences.

To identify the sex chromosomes of *B. zonata*, we applied two new approaches—R-CQ and KAMY—both based on differential read coverage between male and female Illumina libraries (Rallis et al. 2023). Scaffolds 1-7 had similar read coverage in males and females (**Fig. 1B**). Scaffolds 8 and 9 (totaling ∼ 38 Mb) had higher mapping read depth in females than males and were assigned as X-chromosome derived (**Fig. 1B**). An additional ∼4 Mb of putative X-linked sequences was identified in smaller scaffolds. Conversely, scaffolds 10 and 16 (comprising ∼9.2 Mb) showed higher coverage in males and were assigned to the Y chromosome (**Fig. 1B**), with another ∼7.5 Mb of putative Y-linked sequences found on smaller scaffolds. Repeat masking revealed that 81.6% of the X- and 82.13% of the Y-derived scaffolds consisted of repetitive elements, far exceeding the genome-wide average of 55.1% and consistent with observations in other species for the highly-repetitive nature of sex chromosome content (Kaiser and Bachtrog 2010; Bachtrog 2013). The annotation of individual repeat families within the sex chromosome-derived scaffolds suggested similar repeat composition, with LINE retroelements being the most abundant family (X: 22.45%; Y:14.37%).

Mapping WGS reads from both sexes to the two main Y-derived scaffolds (10 and 16) confirmed overall higher coverage in males, although repeat-rich regions attracted female reads as well, supporting shared repetitive content between sex chromosomes and/or autosomes (**Fig. 2A**). To validate Y-linkage of scaffolds 10 and 16, we designed PCR primers to amplify seven unique locations along each scaffold (**Fig. 2A**). All amplicons yielded male-specific products (**Fig. 2B**). We found that scaffold 16 harbors the master-sex determining *MoY -* a gene conserved among Tephritids including all tested *Bactrocera* species (Meccariello et al. 2019; Wu et al. 2024; Fan et al. 2023; Liu et al. 2022) - with its Y-linkage confirmed by PCR (**Fig. 2C**). We identified thirteen copies of *MoY* arranged in head-to-tail tandem repeats; six copies (*BzMoY 6-11*) were identical to the consensus, while the remaining seven exhibited various polymorphisms, including synonymous and non-synonymous substitutions or truncations due to deletions (**Fig. 2D, F**). Consistent with its Y-specific nature, mapping WGS reads to the *MoY*-containing region on scaffold 16 (approximately 0.21–0.29 Mbp) produced male-only alignments over the *MoY* coordinates (**Fig. 2E**). Overall, the identification and validation of Y-chromosome derived sequences in *B. zonata* is expected to support the development of GSS in this species, including through the discovery of unique, Y-linked Cas9-targetable sequences to integrate selectable markers by homology-directed repair linking them to maleness.

**Figure 2.**
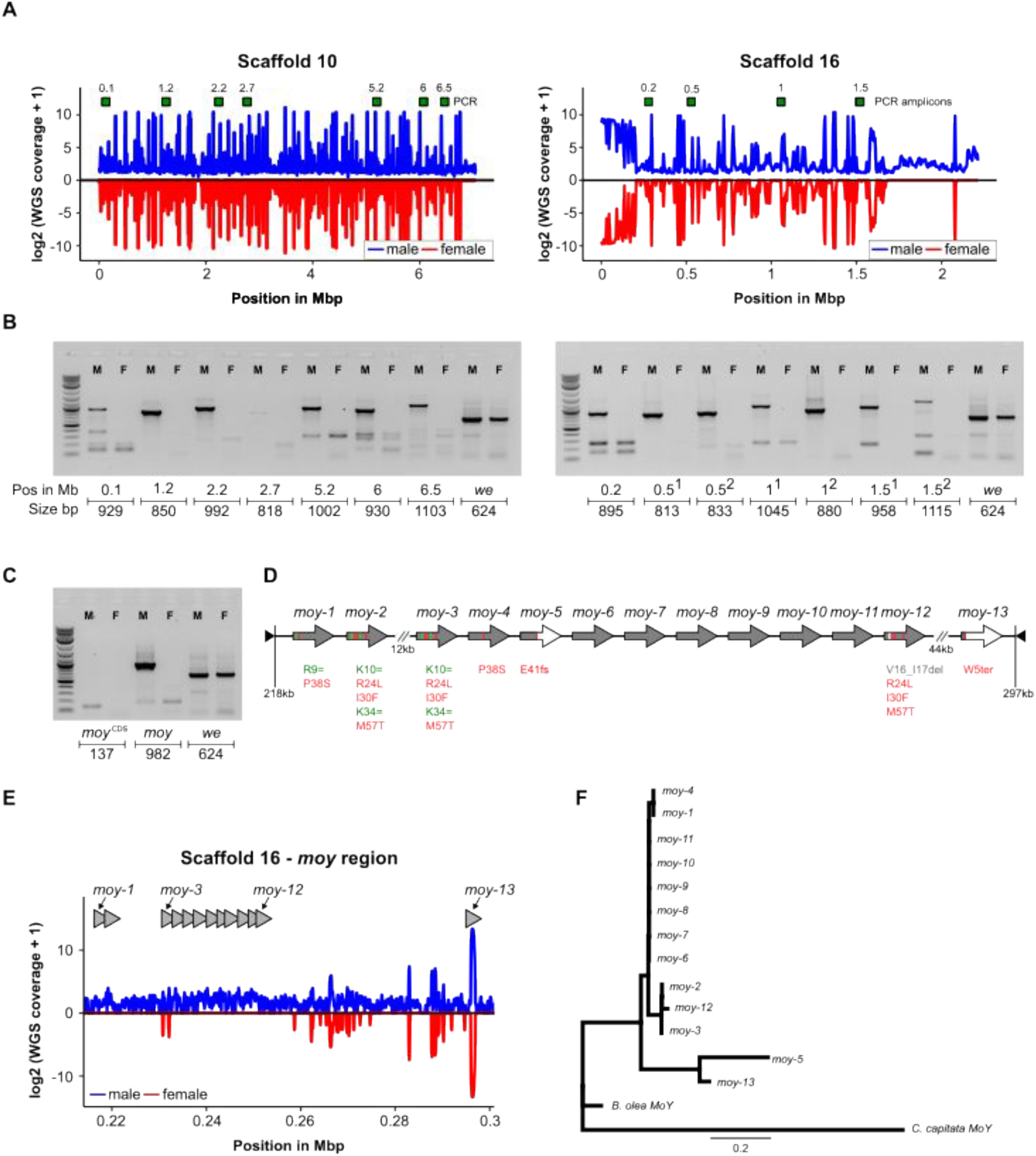
Validation of Y-linked scaffolds and the sex-determining *BzMoY* genes. **A)** Mean coverage from male and female whole-genome sequencing on Y-derived scaffolds 10 and 16 shown in 5 kb sliding windows. Coverage is log2 transformed and female values are displayed on the negative axis for ease of comparison. Green boxes indicate the positions of PCR amplicons, numbered per scaffold. **B)** PCR validation of male-specificity for scaffolds 10 and 16. Amplicons from male (M) and female (F) genomic DNA are shown with size (in bp) and genomic position (in Mbp) indicated. The autosomal *white-eye* gene is used as a positive genomic control in each gel, resulting in PCR amplicons in both sexes. **C)** PCR validation for *BzMoY* - the master sex-determination gene. Two primer pairs were tested, one amplifying a short 137bp *MoY* coding sequence and a longer 982bp fragment including flanking sequences. **D)** Schematic of 13 tandemly repeated *MoY* copies. Six copies (*BzMoY 6–11*) are identical to the consensus and seven others contain various combinations of sequence variants, which are depicted according to HGVS Recommendations (den Dunnen et al. 2016). **E)** Male-specific mapping of WGS reads over the *BzMoY* region on scaffold 16 (0.21–0.29 Mbp). Given smaller region size, mean coverage was calculated in 5o bp sliding windows. **F)** Phylogenetic tree of the 13 *BzMoY* amino acid sequences with validated *MoYs* from *B. olea* and *C. capitata*.

As a first step in identifying selectable markers, we used the *B. tryoni* white-eye protein sequence (Choo et al. 2018) as a query for reciprocal tBLASTn searches against the *B. zonata* genome assembly. A single ortholog was identified on scaffold 3 (34,350,949 - 34,370,584) and its gene model was manually re-annotated (**Fig. 3A**). Its coding sequence was used to construct a phylogenetic tree with other Tephritidae white-eye proteins (**Fig. 3B**), whose topology generally conformed to the Tephritidae phylogeny. White-eye proteins were highly conserved, exhibiting an overall similarity of 96.2% across all species, and 99.7% between *B. zonata* and *B. tryoni*. To confirm that the candidate *B. zonata white-eye* gene is the real ortholog of the functionally validated *B. tryoni white-eye* gene, we performed synteny analysis of flaking sequences (**Fig. 3C**). Synteny of *white-eye* was conserved both upstream and downstream for at least three genes in the *Bactrocera* subgenus, confirming that our selected ortholog is the actual *B. zonata white-eye* gene.

**Figure 3.**
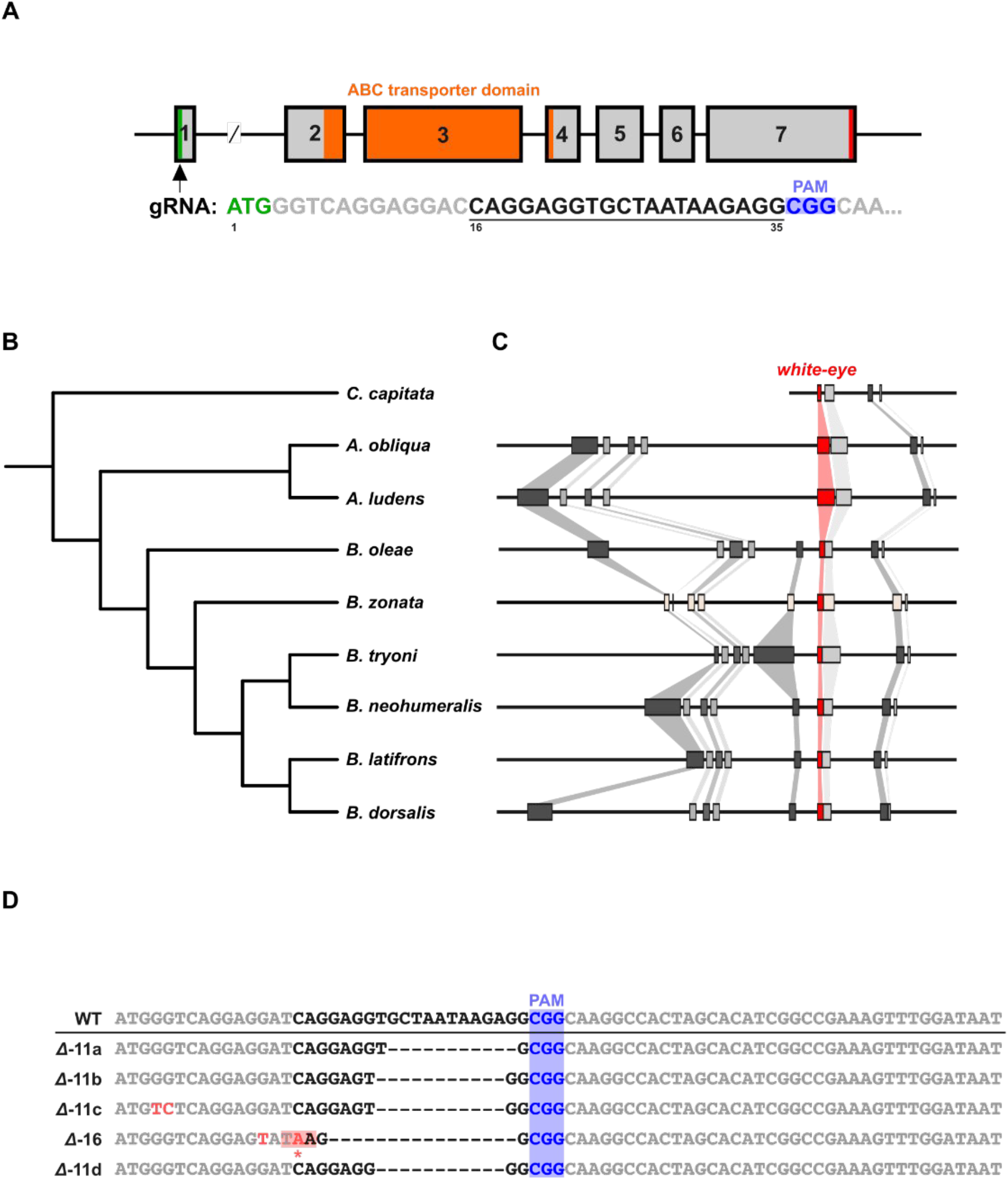
CRISPRing the *white-eye* gene in *B. zonata*. **A)** Gene structure of the *white-eye* locus on scaffold 3, showing gRNA1 target site 16 bp downstream of the start codon. **B)** Phylogenetic tree of White-eye protein sequences from Tephritidae species confirms orthology and sequence conservation. **C)** Conserved synteny surrounding *white-eye* among *Bactrocera* species supports orthology. **D)** Sanger sequencing of G2 white eyed mutants reveals multiple CRISPR-induced indels. The *weΔ-11d* allele contains an 11-bp deletion resulting in a frameshift mutation and is the allele in our homozygous *white-eye* strain.

A guide RNA (gRNA) targeting the first exon - 16 bp downstream of the start codon - was selected (**Fig. 3A**), with no predicted off-targets in the *B. zonata* genome assembly. To avoid using a gRNA targeting polymorphic sites, we sequenced part of the first exon from multiple individuals of our laboratory strain and detected no SNPs over the selected target site. Cas9/gRNA complexes were assembled *in vitro* and injected into syncytial embryos after adapting the medfly microinjection protocol to *B. zonata* in a few important ways (for details, see Material and Methods and Appendix 1). Briefly, dechorionated eggs were aligned for microinjections on a 0.9% agarose platform featuring a vertical 1 mm step, against which eggs were aligned (**Fig. S1**). Injected eggs were incubated on agarose, which provided a moist environment to prevent dehydration during both microinjection and post-injection development. Hatched larvae burrowed into the substrate, remaining viable for several hours, allowing for easier counting and screening. We found that a concentration of 0.9% agarose provided an optimal balance between structural rigidity and optical clarity, facilitating precise injections on rigs using an inverted microscope (**Fig. S1B-C**).

In total, 555 eggs were injected using Cas9 and the *white-eye* gRNA. 188 larvae hatched (33% hatching rate), of which 77 survived to adulthood (41% survival rate from larvae). Among the G0 adults, seven exhibited eye color mosaicism (∼10% mosaicism efficiency) and were individually backcrossed to wild-type (WT) flies. Live genotyping of 20 randomly-selected F1 individuals identified 14 carrying mutant alleles, which were intercrossed to produce homozygous F2 individuals. 403 F2 adults were screened for eye phenotypes, out of which 102 flies (25%) exhibited a white eye phenotype (**Table 2**), with Sanger sequencing confirming the presence of multiple null alleles (**Fig. 3D**).

To isolate a pure strain containing a single mutant allele, white eyed F2 adults were individually backcrossed to WT flies. From six backcross cages, one successfully produced F3 progeny that, when intercrossed, yielded F4 homozygous mutants. The resulting strain, designated *weΔ-11d*, has been maintained for six generations. The white eye phenotype in this strain is due to an 11-bp deletion at position +22 of the first exon, resulting in a frameshift and a nonfunctional protein (**Fig. 3D**). Mutant adults display pearl-white eyes, in contrast to the dark, metallic grey eyes of WT flies (**Fig. 4**). In addition to the eyes, subtle pigmentation differences were also observed in other head structures, including the antero-medial hump, lunule, fronto-orbital bristles, genal spot, postgena, the ocellar triangle and the ocelli. The pair of facial spots in the antennal groove that play an important taxonomic role in species identification are not affected by the *white-eye* mutation (**Fig. 4**).

**Figure 4.**
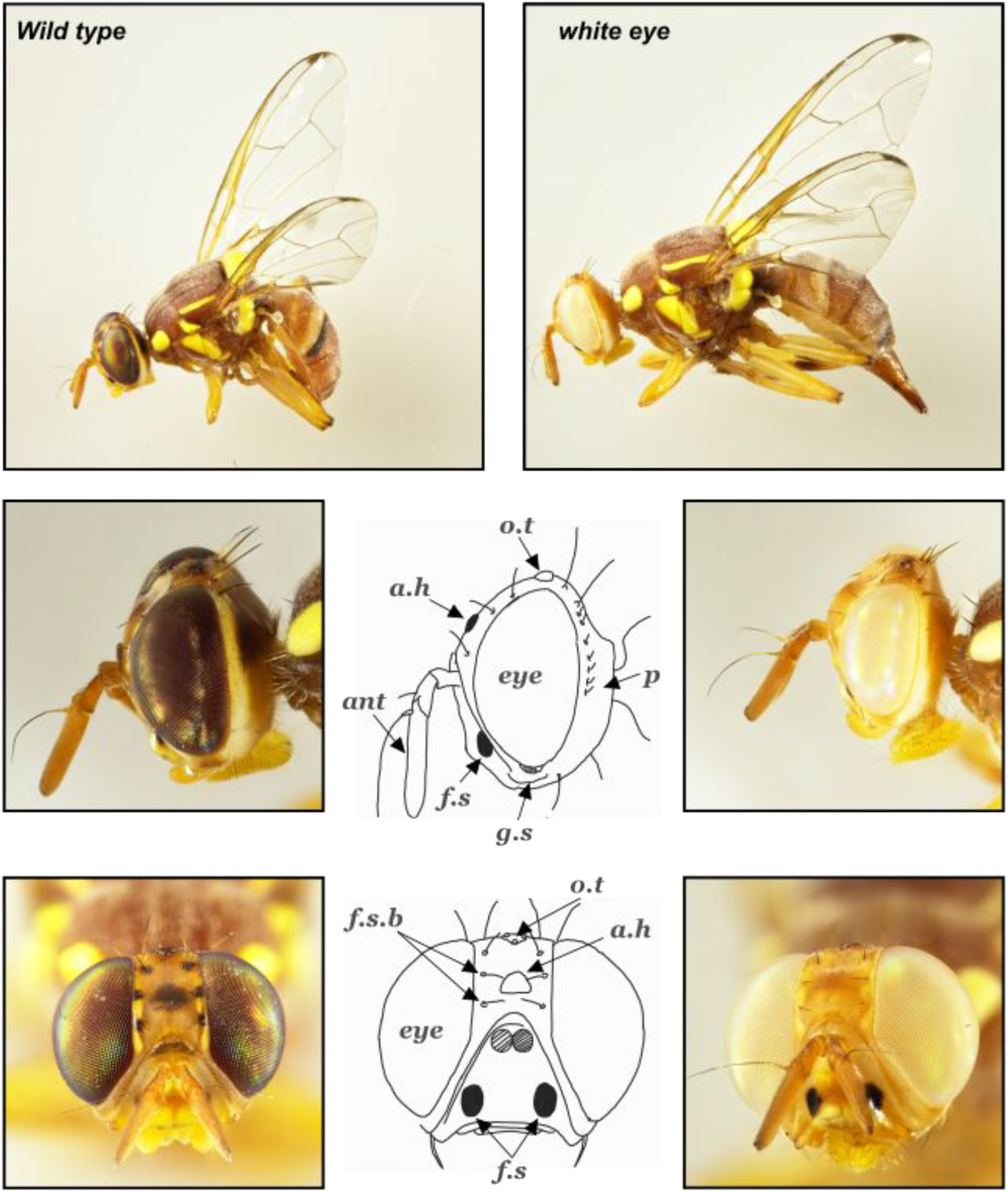
Phenotypic characterization of *white-eye* mutants. Adult *B. zonata* flies homozygous for the *weΔ-11d* allele show a distinct pearl-white eye phenotype compared to the metallic grey eyes of wild-type flies. Additional pigmentation differences are observed in head structures including the antero-medial hump (AH), lunule (L), fronto-orbital bristles (fOB), genal spot (GS), postgena (P), the ocellar triangle (OT) and the ocelli (O). Notably, the facial spots (FS) in the antennal grooves remain unaffected, preserving their diagnostic value for taxonomy.

## Discussion

In the present study, we significantly advance genomic resources and gene-editing methodologies aimed at the long-term development of Genetic Sexing Strains (GSSs) in the invasive fruit fly pest, *Bactrocera zonata*. We report a chromosome-level genome assembly for *B. zonata*, comprising approximately 672 Mbp, providing a robust foundation for genetic and functional studies. This genome assembly contributes to the rapidly growing genomic resources for tephritid fruit flies, alongside recent high-quality genome assemblies for species such as *Anastrepha obliqua* (Sim et al. 2024), *Anastrepha ludens (Congrains et al. 2024), Merzomyia westermanni* (Falk et al. 2024), *Neoceratitis asiatica* (Guo et al. 2023), *Ceratitis capitata, C. quilicii C. rosa, Zeugodacus cucurbitae* (Deschepper et al. 2024; Sim and Geib 2017), and several Bactrocera species, including *B. dorsalis* (Jiang et al. 2022; Carraretto et al. 2020), *B. oleae* (Bayega et al. 2020), *B. zonata* (Deschepper et al. 2024) and *Zeugodacus tau* (Wang et al. 2023). Collectively, these resources represent invaluable data for comparative studies into the biology, ecology and evolution of this important family of economically important pests. They may also provide critical datasets for advancing genetic control strategies, for example by mapping orthologs of important target genes relevant for selection and Y-chromosome sequences to then efficiently select for sex.

Our chromosome-level genome assembly of *B. zonata* successfully resolved autosomes into one or two scaffolds each, which were highly syntenic with *C. capitata* chromosomes. Using differential read coverage methods (KAMY and R-CQ; Rallis et al. 2023), we assigned scaffolds 10 and 16 (comprising ∼9.2 Mbp) as Y chromosome-derived. Both KAMY and R-CQ are based on differential read coverage between male and female Illumina whole-genome sequence libraries and are, essentially, modifications of previous pipelines like CQ (Hall et al. 2013) and YGS (Carvalho and Clark 2013). Subsequent PCR validation confirmed the male specificity of these sequences. Within these scaffolds, we characterized an abundance (82%) of repetitive elements, predominantly LINE retroelements, paralleling observations on the X chromosome. Furthermore, detailed characterization of scaffold 16 revealed a tandemly repeated cluster of the male-determining *BzMoY* gene. These findings substantially enrich our current understanding of Tephritid Y chromosome composition and structure and offer specific genomic targets for future genetic modifications for developing effective GSSs. Moreover, resolving Y chromosome sequences across various tephritid species, as demonstrated here and in recent studies (Bayega et al. 2020; Wu et al. 2024; Meccariello et al. 2019; Liu et al. 2022) enhances comparative studies on Y chromosome evolution and sex determination mechanisms within the Tephritidae.

Beyond genomic advancements, an important technical challenge addressed in this study was the optimization of embryo microinjection protocols. Using agarose as an alignment substrate, we substantially improved egg handling, post-injection survival, and overall injection success. The agarose surface maintained a consistently moist environment, which proved useful for egg viability and optimal embryonic development. Importantly, larvae hatched from injected eggs exhibited natural burrowing behaviors into the agarose, reducing neonatal mortality rates that we observed with alternative substrates such as oil or standard petri-dish surfaces. Additionally, the agarose method allowed for greater flexibility and precision when counting, screening, and transferring hatched larvae, making the procedure both efficient and practical for genetic screening. This optimized protocol has broad applicability and could be effectively generalized to other insect species, particularly within the Tephritidae family. A detailed protocol outlining this optimized method is included (see Appendix 1).

Leveraging these optimized techniques, we demonstrated CRISPR/Cas9-mediated genome editing in *B. zonata* by targeting the conserved *white-eye* gene involved in eye pigmentation across many insect taxa, as a proof-of-concept. The successful generation of a stable homozygous mutant strain (*weΔ-11d*) with an easily identifiable phenotype aligns with similar achievements in closely related species such as *C. capitata* (Meccariello et al. 2017), *Zeugodacus cucurbitae* (Paulo et al. 2022), *B. oleae* (Meccariello et al. 2020), *B. dorsalis* (Zhao et al. 2019) and *B. tryoni* (Choo et al. 2018). Future research should prioritize developing additional selectable phenotypes, particularly *temperature-sensitive lethal* (*tsl*) mutations that facilitate efficient embryonic sex separation of males and females. Such mutations have proven most practical in SIT programs for other fruit fly pests, such as the medfly (Franz, Bourtzis, and Cáceres 2021) (Augustinos et al. 2017). Advances in CRISPR-based targeted insertion methods could further streamline the integration of these genetic markers directly onto the Y chromosome.

In conclusion, our study provides foundational genomic resources, improved technical methodologies, and successful proof-of-concept gene editing outcomes essential for the sustainable genetic management of *B. zonata*. As the research community refines and expands these approaches further, we anticipate that they will contribute meaningfully toward environmentally friendly, species-specific pest control solutions, ultimately reducing reliance on insecticides and enhancing agricultural sustainability and food security globally.

## Figure Captions

**Figure S1. Adapted microinjection setup for *B. zonata*. A)** Preparing agarose substrate for alignment and microinjection. **B)** *B. zonata* egg aligned for microinjection on the agarose 10mm step viewed through a dissecting stereomicroscope. **C)** *B. zonata* egg microinjection viewed through an inverted microscope (i.e through agarose).

## Acknowledgements

We would like to thank Haig Djambazian, Ioannis Ragoussis and Alistair Darby for their work in developing high-quality Tephritids genomics data. We are also thankful to Katerina Nikolouli, Maria-Eleni Grigoriou and Kostas Bourtzis for sharing their protocols and data towards *Bactrocera* CRISPR editing. We thank Marc Schetelig for coordinating efforts in Tephritid GSS development. This study benefited from discussions at meetings for the Coordinated Research Project D44003 on the “Generic approach for the development of genetic sexing strains for SIT applications”, funded by the International Atomic Energy Agency (IAEA).

## Ethics

All insects were handled in accordance with and under the supervision of the ARO Institutional Animal Care and Use Committee approval number 2307-118-2-VOL-IL. All insect work was performed in facilities maintaining Arthropod Containment Level 2. This work received Institutional Approval and relevant authorizations from the Israel Plant Protection and Inspection Services, the Ministry of Agriculture and Food Security (Permit Number 103224 and 1035/21).

## Appendix 1

Microinjection Protocol for *Bactrocera zonata*

1. Egg Collection
  1. Prepare Collection Bottles:
    ∘ Obtain 100 ml commercial yogurt bottles (e.g., Actimel).
    ∘ Sterilize a needle and use it to pierce multiple small holes in the bottle’s surface.
  2. Set Up for Egg Laying:
    ∘ Place a 50 ml Falcon tube, also pierced with small holes, inside the yogurt bottle.
    ∘ Add a small piece of citrus fruit (e.g., orange) to the Falcon tube to attract female flies.
  3. Collect Eggs:
    ∘ Place the prepared bottles in cages with adult *Bactrocera zonata* for 0–2 hours.
    ∘ After the collection period, remove the eggs by gently washing the contents of the bottles with distilled water.
  4. Transfer Eggs:
    ∘ Transfer the eggs into Drosophila egg collection baskets for further processing.
2. Dechorionation
  1. Prepare Dechorionation Solution:
    ∘ Prepare a 50% bleach solution for dechorionation.
  2. Softening of egg chorion:
    ∘ Submerge the eggs in the 50% bleach solution for 25 seconds to soften the chorion layer.
    ∘ Rinse the eggs under running double-distilled water (ddw) for one minute to remove any residual bleach.
3. Agarose Staircase Preparation
  1. Prepare Agarose Solution:
    ∘ Melt 0.9 g of agarose in 100 ml of double-distilled water using a microwave with three pulses of 30 seconds each.
  2. Solidify Agarose:
    ∘ Pour 15 ml of the melted agarose solution onto a microscope glass slide placed in the center of a medium Petri dish.
    ∘ Allow the agarose to solidify for 20 minutes.
  3. Prepare Agarose Mold:
    ∘ Trim the edges of the solidified agarose, leaving a 0.5 cm border around the slide.
    ∘ Remove the slide from the mold and cut the agarose into stair-shaped pieces for eggs alignment.
4. Egg Alignment
  1. Align Eggs:
    ∘ Arrange the dechorionated eggs on the agarose staircase with their dorsal side facing outward and head-to-tail orientation.
    ∘ Position approximately 20 eggs per step of the staircase.
5. Needle Preparation
  1. Prepare Injection Needles:
    ∘ Pull quartz glass capillaries (1 mm outer diameter, 0.70 mm inner diameter) using a Sutter P-2000 Laser Puller.
    ∘ Set the following parameters: Heat 750, Filament 4, Pull 165, Velocity 40, Delay 150.
6. Microinjection
  1. Perform Injections:
    ∘ Inject eggs 2–4 hours after egg laying (AEL) using the prepared quartz glass capillaries.
  2. Post-Injection Incubation:
    ∘ After injection, transfer each staircase to a small Petri dish.
    ∘ Place the Petri dishes in a box containing approximately 3 cm of water to maintain a moist environment for the developing eggs.
    ∘ Incubate the eggs at 26°C for 48 hours.
7. Larval Transfer
  1. Transfer Larvae:
    ∘ Once hatched, transfer the G0 larvae to 12-well plates containing G0 larval food.
    ∘ Each well should accommodate up to 5 larvae.
  2. Provide Suitable Environment:
    ∘ Place the plates in a box with 0.5 cm of ground vermiculite to allow the larvae to burrow and pupate.

## References

Altschul, S. F., W. Gish, W. Miller, E. W. Myers, and D. J. Lipman. 1990. “Basic Local Alignment Search Tool.” Journal of Molecular Biology 215 (3): 403–10.

ArimaGenomics. 2023. Arima Genomics Mapping Pipeline. https://github.com/ArimaGenomics/mapping_pipeline/tree/master.

Augustinos, A. A., A. Targovska, E. Cancio-Martinez, E. Schorn, G. Franz, C. Cáceres, A. Zacharopoulou, and K. Bourtzis. 2017. “Ceratitis Capitata Genetic Sexing Strains: Laboratory Evaluation of Strains from Mass-rearing Facilities Worldwide.” Entomologia Experimentalis et Applicata 164 (3): 305–17.

Bachtrog, Doris. 2013. “Y-Chromosome Evolution: Emerging Insights into Processes of Y-Chromosome Degeneration.” Nature Reviews. Genetics 14 (2): 113–24.

Bayega, Anthony, Haig Djambazian, Konstantina T. Tsoumani, Maria-Eleni Gregoriou, Efthimia Sagri, Eleni Drosopoulou, Penelope Mavragani-Tsipidou, et al. 2020. “De Novo Assembly of the Olive Fruit Fly (Bactrocera Oleae) Genome with Linked-Reads and Long-Read Technologies Minimizes Gaps and Provides Exceptional Y Chromosome Assembly.” BMC Genomics 21 (1): 259.

Carraretto, Davide, Nidchaya Aketarawong, Alessandro Di Cosimo, Mosè Manni Francesca Scolari, Federica Valerio, Anna R. Malacrida, Ludvik M. Gomulski, and Giuliano Gasperi. 2020. “Transcribed Sex-Specific Markers on the Y Chromosome of the Oriental Fruit Fly, Bactrocera Dorsalis.” BMC Genetics 21 (Suppl 2): 125.

Carvalho, Antonio Bernardo, and Andrew G. Clark. 2013. “Efficient Identification of Y Chromosome Sequences in the Human and Drosophila Genomes.” Genome Research 23 (11): 1894–1907.

Choo, A., P. Crisp, R. Saint, L. V. O’Keefe, and S. W. Baxter. 2018. “CRISPR/Cas9-mediated Mutagenesis of the white Gene in the Tephritid Pest Bactrocera Tryoni.” Zeitschrift Für Angewandte Entomologie [Journal of Applied Entomology] 142 (1-2): 52–58.

Congrains, Carlos, Sheina B. Sim, Daniel F. Paulo, Renee L. Corpuz, Angela N. Kauwe, Tyler J. Simmonds, Sheron A. Simpson, Brian E. Scheffler, and Scott M. Geib. 2024. “Chromosome-Scale Genome of the Polyphagous Pest Anastrepha Ludens (Diptera: Tephritidae) Provides Insights on Sex Chromosome Evolution in Anastrepha.” G3 (Bethesda, Md.) 14 (12). 10.1093/g3journal/jkae239.

Deschepper, Pablo, Sam Vanbergen, Lore Esselens, John S. Terblanche, Minette Karsten, Maxi Snyman, Domingos Cugala, et al. 2024. “A New Genome Sequence Resource for Five Invasive Fruit Flies of Agricultural Concern: Ceratitis Capitata, C. Quilicii, C. Rosa, Zeugodacus Cucurbitae and Bactrocera Zonata (Diptera, Tephritidae).” F1000Research 13 (1492): 1492.

Dunnen, Johan T. den, Raymond Dalgleish, Donna R. Maglott, Reece K. Hart, Marc S. Greenblatt, Jean McGowan-Jordan, Anne-Francoise Roux, Timothy Smith, Stylianos E. Antonarakis, and Peter E.M. Taschner. 2016. “HGVS Recommendations for the Description of Sequence Variants: 2016 Update.” Human Mutation 37 (6): 564–69.

El-Gendy, I. 2022. “Bactrocera Zonata (peach Fruit Fly).” CABI Compendium CABI Compendium (January). 10.1079/cabicompendium.17694.

Ewart, G. D., and J. Howells. 1998. “ABC Transporters Involved in Trans-Port of Eye Pigment Precursors in Drosophila Melanogaster.” Methods in Enzymology 292:213–24.

Falk, Steven, Liam M. Crowley, Nathan C. Medd, University of Oxford and Wytham Woods Genome Acquisition Lab, Darwin Tree of Life Barcoding collective, Wellcome Sanger Institute Tree of Life Management, Samples and Laboratory team, Wellcome Sanger Institute Scientific Operations: Sequencing Operations, Wellcome Sanger Institute Tree of Life Core Informatics team, Tree of Life Core Informatics collective, and Darwin Tree of Life Consortium. 2024. “The Genome Sequence of a Tephritid Fruit Fly, Merzomyia Westermanni Meigen 1826.” Wellcome Open Research 9 (August):480.

Fan, Zi-Zhen, Qin Ma, Si-Ya Ma, Feng-Qin Cao, Ri-Hui Yan, and Xian-Wu Lin. 2023. “Maleness-on-the-Y (MoY) Orthologue Is a Key Regulator of Male Sex Determination in Zeugodacus Cucurbitae (Diptera: Tephritidae).” Journal of Integrative Agriculture 22 (2): 505–13.

Flynn, Jullien M., Robert Hubley, Clément Goubert, Jeb Rosen, Andrew G. Clark, Cédric Feschotte, and Arian F. Smit. 2020. “RepeatModeler2 for Automated Genomic Discovery of Transposable Element Families.” Proceedings of the National Academy of Sciences of the United States of America 117 (17): 9451–57.

Franz, G., K. Bourtzis, and C. Cáceres. 2021. “Practical and Operational Genetic Sexing Systems Based on Classical Genetic Approaches in Fruit Flies, an Example for Other Species Amenable to Large-Scale Rearing for the Sterile Insect Technique Chapter 4.3.” In Sterile Insect Technique, 575–604. CRC Press.

Gazit, Yoav, and Ruti Akiva. 2017. “Toxicity of Malathion and Spinosad to Bactrocera Zonata and Ceratitis Capitata (Diptera: Tephritidae).” The Florida Entomologist 100 (2): 385–89.

Guan, Dengfeng, Shane A. McCarthy, Jonathan Wood, Kerstin Howe, Yadong Wang, and Richard Durbin. 2020. “Identifying and Removing Haplotypic Duplication in Primary Genome Assemblies.” Bioinformatics (Oxford, England) 36 (9): 2896–98.

Guo, Shaokun, Bo Liu, Jia He, Zihua Zhao, Rong Zhang, and Zhihong Li. 2023. “Chromosome-Level Genome Assembly of an Important Wolfberry Fruit Fly (Neoceratitis Asiatica Becker).” Scientific Data 10 (1): 675.

Hall, Andrew Brantley, Yumin Qi, Vladimir Timoshevskiy, Maria V. Sharakhova, Igor V. Sharakhov, and Zhijian Tu. 2013. “Six Novel Y Chromosome Genes in Anopheles Mosquitoes Discovered by Independently Sequencing Males and Females.” BMC Genomics 14 (April):273.

Jiang, Fan, Liang Liang, Jing Wang, and Shuifang Zhu. 2022. “Chromosome-Level Genome Assembly of Bactrocera Dorsalis Reveals Its Adaptation and Invasion Mechanisms.” Communications Biology 5 (1): 25.

Kaiser, Vera B., and Doris Bachtrog. 2010. “Evolution of Sex Chromosomes in Insects.” Annual Review of Genetics 44 (1): 91–112.

Kolmogorov, Mikhail, Jeffrey Yuan, Yu Lin, and Pavel A. Pevzner. 2019. “Assembly of Long, Error-Prone Reads Using Repeat Graphs.” Nature Biotechnology 37 (5): 540–46.

Langmead, Ben, Cole Trapnell, Mihai Pop, and Steven L. Salzberg. 2009. “Ultrafast and Memory-Efficient Alignment of Short DNA Sequences to the Human Genome.” Genome Biology 10 (3): R25.

Li, Heng. 2018. “Minimap2: Pairwise Alignment for Nucleotide Sequences.” Bioinformatics (Oxford, England) 34 (18): 3094–3100.

Li, Heng, and Richard Durbin. 2009. “Fast and Accurate Short Read Alignment with Burrows-Wheeler Transform.” Bioinformatics (Oxford, England) 25 (14): 1754–60.

Liu, Peipei, Wenping Zheng, Jiao Qiao, Ziniu Li, Zhurong Deng, Yimin Yuan, and Hongyu Zhang. 2022. “Early Embryonic Transcriptomes of Zeugodacus Tau Provide Insight into Sex Determination and Differentiation Genes.” Insect Science 29 (3): 915–31.

Mak, Q. X. Charles, Ryan R. Wick, James Matthew Holt, and Jeremy R. Wang. 2023. “Polishing DE Novo Nanopore Assemblies of Bacteria and Eukaryotes with FMLRC2.” Molecular Biology and Evolution 40 (3): msad048.

Meccariello, Angela, Simona Maria Monti, Alessandra Romanelli, Rita Colonna, Pasquale Primo, Maria Grazia Inghilterra, Giuseppe Del Corsano, et al. 2017. “Highly Efficient DNA-Free Gene Disruption in the Agricultural Pest Ceratitis Capitata by CRISPR-Cas9 Ribonucleoprotein Complexes.” Scientific Reports 7 (1): 10061.

Meccariello, Angela, Marco Salvemini, Pasquale Primo, Brantley Hall, Panagiota Koskinioti, Martina Dalíková, Andrea Gravina, et al. 2019. “Maleness-on-the-Y (MoY) Orchestrates Male Sex Determination in Major Agricultural Fruit Fly Pests.” Science (New York, N.Y.) 365 (6460): 1457–60.

Meccariello, Angela, Konstantina T. Tsoumani, Andrea Gravina, Pasquale Primo, Martina Buonanno, Kostas D. Mathiopoulos, and Giuseppe Saccone. 2020. “Targeted Somatic Mutagenesis through CRISPR/Cas9 Ribonucleoprotein Complexes in the Olive Fruit Fly, Bactrocera Oleae.” Archives of Insect Biochemistry and Physiology 104 (2): e21667.

Musapa, Mulenga, Taida Kumwenda, Mtawa Mkulama, Sandra Chishimba, Douglas E. Norris, Philip E. Thuma, and Sungano Mharakurwa. 2013. “A Simple Chelex Protocol for DNA Extraction from Anopheles Spp.” Journal of Visualized Experiments: JoVE, no. 71 (January). 10.3791/3281.

Palmer, Jonathan M., and Jason Stajich. 2020. Funannotate v1.8.1: Eukaryotic Genome Annotation. Zenodo. 10.5281/ZENODO.1134477.

Paulo, Daniel F., Alex Y. Cha, Angela N. Kauwe, Keena Curbelo, Renee L. Corpuz, Tyler J. Simmonds, Sheina B. Sim, and Scott M. Geib. 2022. “A Unified Protocol for CRISPR/Cas9-Mediated Gene Knockout in Tephritid Fruit Flies Led to the Recreation of White Eye and White Puparium Phenotypes in the Melon Fly.” Journal of Economic Entomology 115 (6): 2110–15.

Perez, Gerardo, Galt P. Barber, Anna Benet-Pages, Jonathan Casper, Hiram Clawson, Mark Diekhans, Clay Fischer, et al. 2025. “The UCSC Genome Browser Database: 2025 Update.” Nucleic Acids Research 53 (D1): D1243–49.

Quinlan, Aaron R., and Ira M. Hall. 2010. “BEDTools: A Flexible Suite of Utilities for Comparing Genomic Features.” Bioinformatics (Oxford, England) 26 (6): 841–42.

Rallis, Dimitris, Konstantina T. Tsoumani, Flavia Krsticevic, Philippos Aris Papathanos, Kostas D. Mathiopoulos, and Alexie Papanicolaou. 2023. “Revisiting Y-Chromosome Detection Methods: R-CQ and KAMY Efficiently Identify Y Chromosome Sequences in Tephritidae Insect Pests.” bioRxiv. 10.1101/2023.10.27.564325.

Simão Felipe A., Robert M. Waterhouse, Panagiotis Ioannidis, Evgenia V. Kriventseva, and Evgeny M. Zdobnov. 2015. “BUSCO: Assessing Genome Assembly and Annotation Completeness with Single-Copy Orthologs.” Bioinformatics (Oxford, England) 31 (19): 3210–12.

Sim, Sheina B., Carlos Congrains, Sandra M. Velasco-Cuervo, Renee L. Corpuz, Angela N. Kauwe, Brian Scheffler, and Scott M. Geib. 2024. “Genome Report: Chromosome-Scale Genome Assembly of the West Indian Fruit Fly Anastrepha Obliqua (Diptera: Tephritidae).” G3 (Bethesda, Md.) 14 (4): jkae024.

Sim, Sheina B., and Scott M. Geib. 2017. “A Chromosome-Scale Assembly of the Bactrocera Cucurbitae Genome Provides Insight to the Genetic Basis of White Pupae.” G3 (Bethesda, Md.) 7 (6): 1927–40.

Tarailo-Graovac, Maja, and Nansheng Chen. 2009. “Using RepeatMasker to Identify Repetitive Elements in Genomic Sequences.” Et Al [Current Protocols in Bioinformatics] Chapter 4 (1): 4.10.1–4.10.14.

Venkat, Bandi, and Gutwin Carl. 2020. “Interactive Exploration of Genomic Conservation.” In Graphics Interface. https://graphicsinterface.org/proceedings/gi2020/gi2020-9/.

Walker, Bruce J., Thomas Abeel, Terrance Shea, Margaret Priest, Amr Abouelliel, Sharadha Sakthikumar, Christina A. Cuomo, et al. 2014. “Pilon: An Integrated Tool for Comprehensive Microbial Variant Detection and Genome Assembly Improvement.” PloS One 9 (11): e112963.

Wang, Yi-Ting, Li-Jun Cao, Jin-Cui Chen, Wei Song, Wei-Hua Ma, Jing-Fang Yang, Xu-Yuan Gao, et al. “Chromosome-Level Genome Assembly of an Agricultural Pest Zeugodacus Tau (Diptera: Tephritidae).” Scientific Data 10 (1): 848.

White, I. M., and M. M. Elson-Harris. 1992. Fruit Flies of Economic Significance: Their Identification and Bionomics., (xii + 601 Pp.), CAB International, Fruit Flies of Economic Significance: Their Identification and Bionomics. English, Book, UK, 9780851987903, Wallingford.

Wu, Shuangxiong, Jiahong Wu, Quan Lei, Donghai He, Xinrui Jiang, Chao Ye, Dong Wei, Jinjun Wang, Luohao Xu, and Hongbo Jiang. 2024. “The Assembly of Y Chromosome Reveals Amplification of Genes Regulating Male Fertility in Bactrocera Dorsalis.” Zoology. bioRxiv. https://www.biorxiv.org/content/10.1101/2024.08.01.606120v1.full.

Yesmin, F., M. Uddin, G. Rahman, and Mohammed Hasanuzzaman. 2019. “Identification of Larval Salivary Gland Polytene Chromosomes of the Peach Fruit Fly, Bactrocera Zonata (Saunders) (Diptera: Tephritidae).” Journal of Biological Control, December. 10.18311/jbc/2019/22718.

Zhao, Santao, Zengzhu Xing, Zhonggeng Liu, Yanhui Liu, Xiangrui Liu, Zhe Chen, Jiahui Li, and Rihui Yan. 2019. “Efficient Somatic and Germline Genome Engineering of Bactrocera Dorsalis by the CRISPR/Cas9 System: Genome Engineering of Bactrocera Dorsalis by CRISPR.” Pest Management Science 75 (7): 1921–32.

Zhou, Chenxi, Shane A. McCarthy, and Richard Durbin. 2023. “YaHS: Yet Another Hi-C Scaffolding Tool.” Bioinformatics (Oxford, England) 39 (1): btac808.

Zingore, Kumbirai M., George Sithole, Elfatih M. Abdel-Rahman, Samira A. Mohamed, Sunday Ekesi, Chrysantus M. Tanga, and Mohammed E. E. Mahmoud. 2020. “Global Risk of Invasion by Bactrocera Zonata: Implications on Horticultural Crop Production under Changing Climatic Conditions.” PloS One 15 (12): e0243047.

