## Supplementary material for "CRISPR/Cas9-mediated mutagenesis of the *white-eye* gene in the tephritid pest *Bactrocera zonata*": Fig. S1

### Figure S1

A

**Preparing agarose microinjection step**

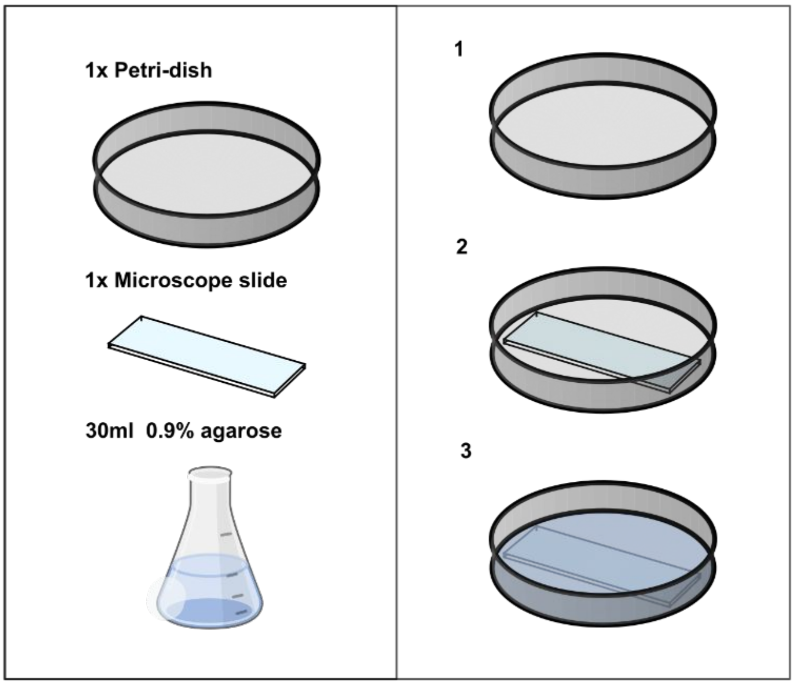


B

**Alignment view (Dissection Microscope)**

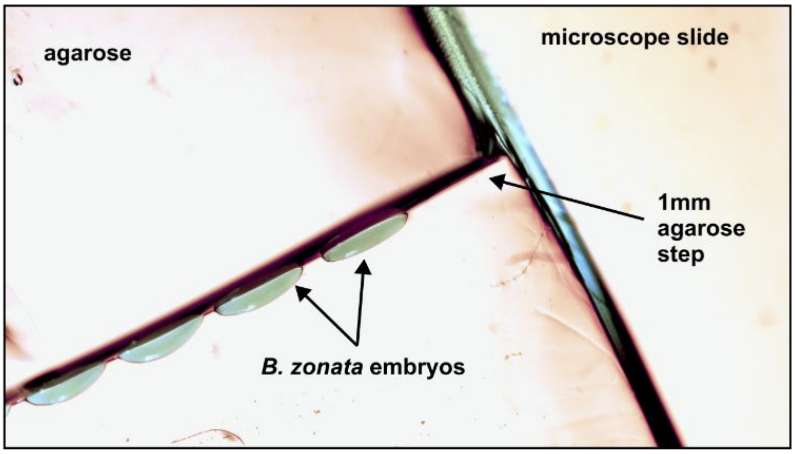


C

**Micro-injection view (Inverted Microscope)**

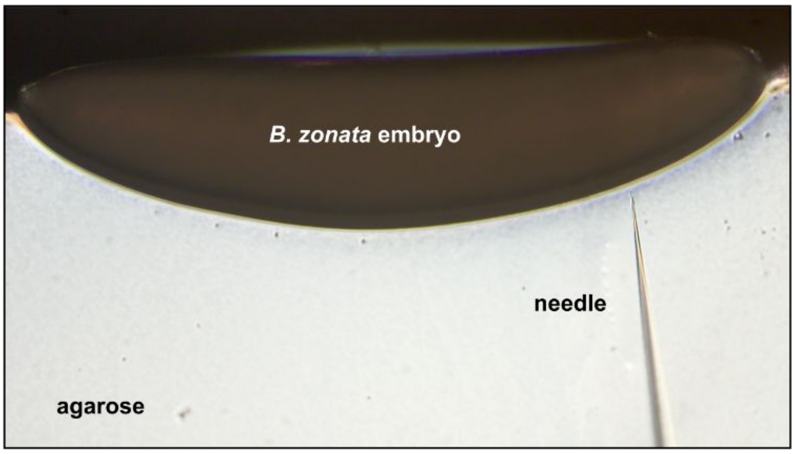
