## Supplementary material for "CRISPR/Cas9-mediated mutagenesis of the *white-eye* gene in the tephritid pest *Bactrocera zonata*": Table 1

**Table 1:** Primers list

| Name | Sequence | Amplicon |
|---|---|---|
| PrimerD_white_ex1_F | GCCCACGCCAATACATTGTTG | white ex1 |
| PrimerE_white_ex1_F | GCATCAATATCCAACCTCCGC | white ex1 |
| PrimerF_white_ex1_R | CTTCAGCTGTCTTTCATTTGCCTC | white ex1 |
| PrimerG_white_ex1_R | CTGTGTCTGTCCATTCGTCGC | white ex1 |
| Scaff_10_0.13M_F | GGTAGAGGAGGGGGTTTTCT | 10_0.13M |
| Scaff_10_0.13M_R | TCCAGGTCTCTACAGGCTTC | 10_0.13M |
| Scaff_10_1.2M_F | ATACTGCGAGGAAGATCCGA | 10_1.2M |
| Scaff_10_1.2M_R | TCCGAGCCACAACTCTTCTA | 10_1.2M |
| Scaff_10_2.2M_F | GGTATACGCGGCATCAAGAT | 10_2.2M |
| Scaff_10_2.2M_R | TTCCGGCGTTAACCTTTGAA | 10_2.2M |
| Scaff_10_2.7M_F | TTTCGAGGTGCGTTGTTACA | 10_2.7M |
| Scaff_10_2.7M_R | CTCTCATCACCAATGCCTCGT | 10_2.7M |
| Scaff_10_5.2M_F | CCATCAACGCCATTCACAAC | 10_5.2M |
| Scaff_10_5.2M_R | GATCAAGAAGCTTACCGGCA | 10_5.2M |
| Scaff_10_6.0M_F | TCAGACAAATCCCAACAGCC | 10_6.0M |
| Scaff_10_6.0M_R | GTAAAATGCCCATTCCGCAC | 10_6.0M |
| Scaff_10_6.5M_F | TGTGTGTCGGTGAAGAGGTA | 10_6.5M |
| Scaff_10_6.5M_R | AAGTGTGGTGACTCGGTAGA | 10_6.5M |
| Scaff_16_0.27M_F | CCTTGCTTGATTAGGCCGAT | 16_0.27M |
| Scaff_16_0.27M_R | GGGCTGTCATAGGTTGATGG | 16_0.27M |
| Scaff_16_0.52M_1_F | ACGCGGTATTCAGATGCATT | 16_0.52M |
| Scaff_16_0.52M_1_R | AACCCGGAAATAGCGACTTC | 16_0.52M |
| Scaff_16_0.52M_2_F | TTGTCTGAAGGTGCAGCAAT | 16_0.52M |
| Scaff_16_0.52M_2_R | TGGCTAGGTTGAAAGATCGC | 16_0.52M |
| Scaff_16_1M_1_F | AGTAGGATGCAAAACGCCAT | 16_1M |
| Scaff_16_1M_1_R | TGAACGCACAGGTTTTACCA | 16_1M |
| Scaff_16_1M_2_F | TTCAGAAGGCCAGATATGCG | 16_1M |
| Scaff_16_1M_2_R | CGGGTCGCATAACATCCTAG | 16_1M |
| Scaff_16_1.5M_1_F | TTGCAAACACTCGGGAATGA | 16_1.5M |
| Scaff_16_1.5M_1_R | ATCCCCTGGCCCTATATTCC | 16_1.5M |
| Scaff_16_1.5M_2_F | CGAATTAGGTCACACCAGGG | 16_1.5M |
| Scaff_16_1.5M_2_R | GTTGATGTGTCAATGCTGCC | 16_1.5M |
| Scaff_16_MoyCDS_F | GTATTCTGTCTAGAAGAATTTGGAATG | 16_MoyCDS |
| Scaff_16_MoyCDS_R | CTTTCGAGTGAACAAACACTTTTTATT | 16_MoyCDS |
| Scaff_16_Moy_F | AGAATTTGTGGAACCTGAGCA | 16_Moy |
| Scaff_16_Moy_R | TTACACAATCCACCCGCAAA | 16_Moy |
