## Supplementary material for "CRISPR/Cas9-mediated mutagenesis of the *white-eye* gene in the tephritid pest *Bactrocera zonata*": Table 2

**Table 2:** 403 F2 adults from three F1 intercross cages were screened for eye phenotypes. 102 flies (25%) exhibited a white eye phenotype.

| Cage | Adults | White eyed adults | White eyed adults (%) |
|---|---|---|---|
| 1 | 152 | 38 | 0.25 |
| 2 | 129 | 34 | 0.264 |
| 3 | 122 | 30 | 0.246 |
| **Total** | 403 | 102 | 0.25 |
